# Black Queen Hypothesis, partial privatization, and quorum sensing evolution

**DOI:** 10.1101/2022.03.11.483843

**Authors:** Lucas Santana Souza, Yasuhiko Irie, Shigetoshi Eda

## Abstract

Microorganisms produce costly cooperative goods whose benefit is partially shared with nonproducers, called ‘mixed’ goods. The Black Queen Hypothesis predicts that partially privatization of benefits from the mixed goods has two major evolutionary implications. First, to favor strains producing several mixed goods over nonproducing strains. Second, to favor the maintenance of cooperative traits through different strains instead of having all cooperative traits present in a single strain (metabolic specialization). Despite the importance of quorum sensing regulation of mixed goods, it is not clear how partial privatization of benefits affects quorum sensing evolution. Here, we studied the influence of partial privatization of benefits on the evolution of quorum sensing. We developed a mathematical population genetics model of an unstructured microbial population considering four strains that differ in their ability to produce an autoinducer (quorum sensing signaling molecule) and a mixed good. Our model assumes that the production of the autoinducers and the mixed goods is constitutive and/or depends on quorum sensing. Our results suggest that partially privatized benefits cannot foster quorum sensing. This result occurs because: (1) a strain that produces both autoinducer and good (fully producing strain) cannot persist in the population; (2) the strain only producing the autoinducer and the strain producing mixed goods in response to the autoinducers cannot coexist, i.e., metabolic specialization cannot be fostered.

## 1. Introduction

The maintenance of microbial strains producing costly beneficial goods (cooperators) is challenging to explain because nonproducing strains (cheaters) are expected to outcompete producing strains [1,2] by taking advantage of the benefits of goods without paying the cost of producing goods. Growing evidence shows that benefits are not always equally shared among producers and nonproducers [3–5]. These goods, providing privatized and public benefits, have been called “mixed” goods [6]. Morris et al. proposed the Black Queen Hypothesis which predicts that the growth of mixed good producers is fostered over that of nonproducers whenever the privatized benefits offset the costs of producing goods [3–5,7].

The production of mixed goods is regulated by the population density-dependent gene regulation of bacteria, called quorum sensing (QS) and/or a QS-independent mechanism. One example of mixed goods is the siderophores in *Pseudomonas aeruginosa* [7–9], which is known to be a virulence factor of the bacteria [10,11]. The benefit of siderophore is partially privatized as only a proportion of the molecules is secreted from the bacteria [7,8]. The partially secreted siderophore provides benefits to nonproducing strains. Another mechanism of partial privatization include the intracellular cleavage of molecules into smaller molecules, some of which are available to the environment [3].

The Black Queen Hypothesis predicts that cooperation can be maintained in two main ways. By favoring a strain producing all cooperative goods (a fully producing strain) [12–14] or by favoring different complementary strains, each producing a cooperative good (i.e., metabolic specialization) [12,13]. These two predictions were also found in models that involved constitutively (QS-independent) production of goods. However, despite growing evidence that QS regulates mixed goods, no model analyzed whether these two predictions are valid for QS evolution.

There are two reasons for why it is unclear whether partially privatized benefits can foster QS via a fully producing strain which produces both QS signaling molecules (autoinducers) and goods. First, the costs of QS-independent production of partially privatized goods are lower than that of QS-regulated production of the goods. Thus, if a mixed good is regulated by a costless, QS-independent mechanism (e.g., constitutively produced), then benefits only need to offset the mixed goods’ costs [3]. However, when mixed goods are QS regulated, privatized benefits need to offset not only the costs of mixed goods but also autoinducers’ cost. Second, experiments that involve QS regulation of a mixed good have successfully shown that the partially privatized benefits suppress the invasion of strains that are not producing mixed goods [15]. The problem is that these studies on the interactions of two strains might not reflect the case when more than two strains are present [16]. Thus, it is not clear whether partially privatized benefits from mixed goods support the growth of fully producing strain.

Additionally, whether partially privatized benefits foster QS via metabolic specialization (i.e., via one strain only producing autoinducers and the other only responding to it) is unclear. That is because metabolic specialization has only been demonstrated by mathematical models assuming a costless regulation of two mixed goods (13,14,17). The problem is that while a QS-independent regulation of two mixed goods might be costless, a QS-regulation requires the production of autoinducers, which is costly. Hence, despite metabolic specialization being a prediction of the Black Queen Hypothesis [12,14], whether QS can be maintained through metabolic specialization needs to be tested.

Here, we built a mathematical model and conducted a theoretical study to examine whether QS can be maintained in the population due to the partial privatization of benefits from a mixed good. Our mathematical model evaluated whether QS is favored by either the maintenance in the population of (A) a strain that produces both autoinducers and mixed goods (fully producing strain); or (B) through the coexistence of two strains, one only producing autoinducers and the other only producing mixed goods (metabolic specialization). Our theoretical study predicted that partially privatized benefits from mixed goods cannot maintain autoinducer-producing and good-producing strains.

## 2. Results

In our model, we assumed that two types of costly molecules are produced, namely an autoinducer (QS signaling molecule) and a mixed good. We considered four strains, each carrying one of two different alleles in two gene loci. At locus ***A***, allele *A* produces an autoinducer that diffuses throughout the environment. Allele *a* is incapable of producing the autoinducer. At locus ***G***, allele *G* recognizes the autoinducers and produces a mixed good through a QS-independent (costless constitutive production) and QS-dependent (i.e., in response to autoinducers frequency) mechanisms. Allele *g* cannot recognize the autoinducers and cannot produce the mixed good.

In nature, mixed goods can provide benefits by reducing cell death [5,7], or promoting cell growth [3]. Here, we assume that the private and public benefits of the mixed good promote growth. The production of the mixed goods and the autoinducers might occur solely by QS-dependent mechanism or together with QS-dependent mechanism. The four strains considered in this work are summarized in the Table 1. The strain *AG* regulates the production of the autoinducer and the mixed good by QS-independent and QS-dependent mechanisms. The strain *aG* cannot produce the autoinducer but produces the mixed good by QS-independent and QS-dependent regulation. The strain *Ag* produces the autoinducer through a QS-independent mechanism but not the mixed good. The strain *ag* neither produces autoinducer nor the mixed good. For simplicity, hereafter, mixed goods will be just referred as goods.

**Table 1.** Strain types and a description of their allele functions

| STRAIN | ALLELE FUNCTION |
| --- | --- |
| *AG* | *Autoinducer-producer/Good-producer* |
| *aG* | *Autoinducer-nonproducer/Good-producer* |
| *Ag* | *Autoinducer-producer/Good-nonproducer* |
| *ag* | *Autoinducer-nonproducer/Good-nonproducer* |

Here, we analyzed the evolution of the population genetics by tracking each strain’s frequency. To evaluate whether partially privatized benefits foster QS, we first checked whether the strain *AG* could be maintained when in pairwise interaction with *aG, Ag*, or *ag*. Since AG can produce and recognize the autoinducer, we consider QS is maintained when AG is present in the population. QS could also be maintained by the coexistence between *Ag* and *aG*, we tested whether selection favors their coexistence (favor metabolic specialization). Lastly, we tested whether partially privatized benefits can favor QS, the maintenance of alleles *A* and *G*, when all strains are simultaneously present in the population.

### (A) Pairwise analysis of *AG* and *ag*

As depicted in Figure 1a, *ag* exploits *AG* by neither producing the autoinducer nor the good. Here, we analyzed whether QS can be maintained in a population by checking if selection fosters pure populations of *AG* or the coexistence of *AG* and *ag*.

**Figure 1.**
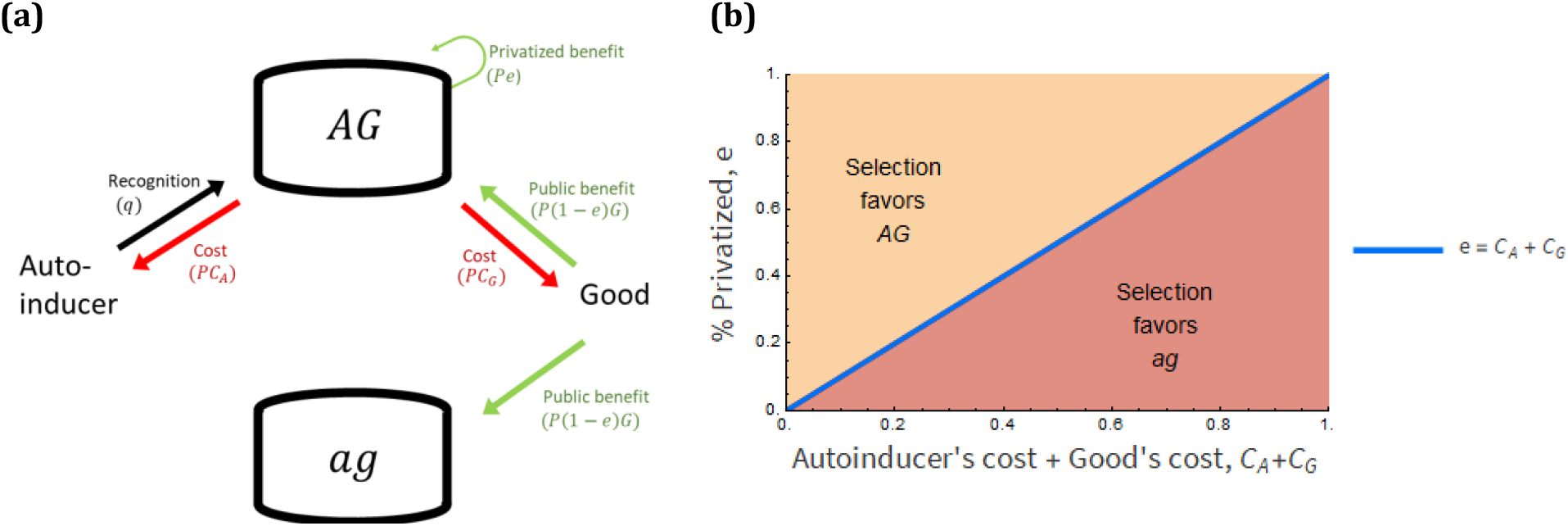
The partially privatized benefit enables *AG* to outcompete *ag*. **(a)** A schematic of the social interaction between strains *AG* and *ag*. Red arrows indicate a costly production of a good and an autoinducer. The black arrow indicates that the autoinducer triggers the production of functional alleles (*A* and *G*). Green arrows indicate the access to the good’s benefits. *AG* accesses both public and private benefits, while *ag* accesses only public benefit. **(b)** A geometric visualization of which strain selection favors. In yellow and red areas, selection favors pure populations of *AG* and *ag*, respectively. The blue line is where both strains are equally fit, which happens when the per capita partially privatized benefit equalizes to the sum of the per capita costs of producing an autoinducer and a good. This result is valid for parameters: 0 ≤ *L* ≤ 1 and having either *q* ≠ 0, or *q*_0_ ≠ 0 or 0 < *q*_0_, *q* ≤ 0.5. Please see Table 2 and Section 6B for the definition of letters (q, C_A_ etc.) and the equations used for the analysis, respectively.

We found that—at any initial ratio of both strains—selection fosters pure populations of *AG* if the per capita partially privatized benefit offsets the per capita cost of producing both the good and the autoinducer. Otherwise, selection favors pure populations of *ag* (Figure 1b). In sum, the factor that determines the outcome of selection is the relative amount of privatized benefit to cost—not the absolute amount of privatized benefit (Section 6.B).

**Table 2.**
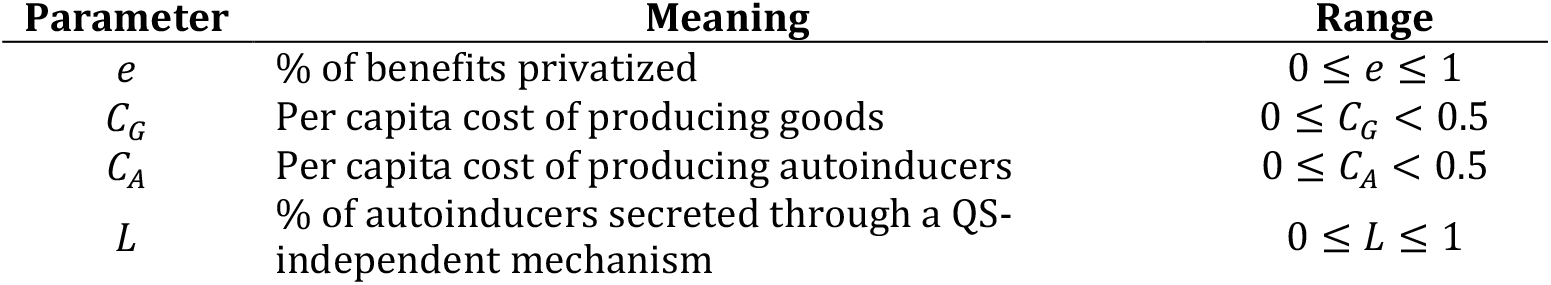
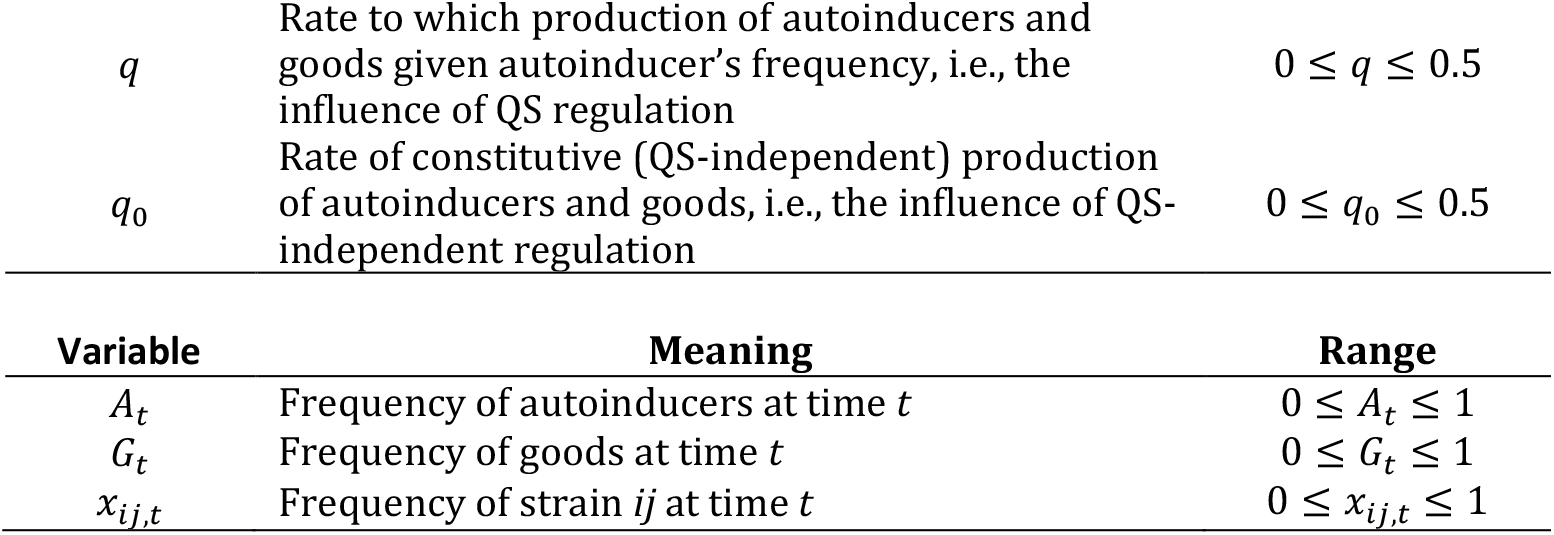
List of parameters and variables and their meaning and range.

| Parameter | Meaning | Range |
| --- | --- | --- |
| $e$ | % of benefits privatized | $0 \leq e \leq 1$ |
| $C_G$ | Per capita cost of producing goods | $0 \leq C_G < 0.5$ |
| $C_A$ | Per capita cost of producing autoinducers | $0 \leq C_A < 0.5$ |
| $L$ | % of autoinducers secreted through a QS-independent mechanism | $0 \leq L \leq 1$ |
| $q$ | Rate to which production of autoinducers and goods given autoinducer's frequency, i.e., the influence of QS regulation | $0 \leq q \leq 0.5$ |
| $q_0$ | Rate of constitutive (QS-independent) production of autoinducers and goods, i.e., the influence of QS-independent regulation | $0 \leq q_0 \leq 0.5$ |

| Variable | Meaning | Range |
| --- | --- | --- |
| $A_t$ | Frequency of autoinducers at time $t$ | $0 \leq A_t \leq 1$ |
| $G_t$ | Frequency of goods at time $t$ | $0 \leq G_t \leq 1$ |
| $x_{ij,t}$ | Frequency of strain $ij$ at time $t$ | $0 \leq x_{ij,t} \leq 1$ |

Moreover, we found that the ability to produce autoinducers and goods in response to autoinducers neither favors nor disfavors any of the two strains (Equation 6). Similarly, the ability to produce autoinducers and goods by a QS-independent mechanism neither favors nor disfavors any of the two strains. The type of regulation, QS-dependent or QS-independent does not matter because only *AG* produces autoinducers and goods.

### (B) Pairwise analysis of *AG* and *Ag*

As depicted in Figure 2A, *Ag* exploits *AG* by not producing the good and by producing fewer autoinducers. Here, we analyzed whether QS can be maintained in a population by checking if selection fosters pure populations of *AG* or the coexistence of *AG* and *Ag*.

**Figure 2.**
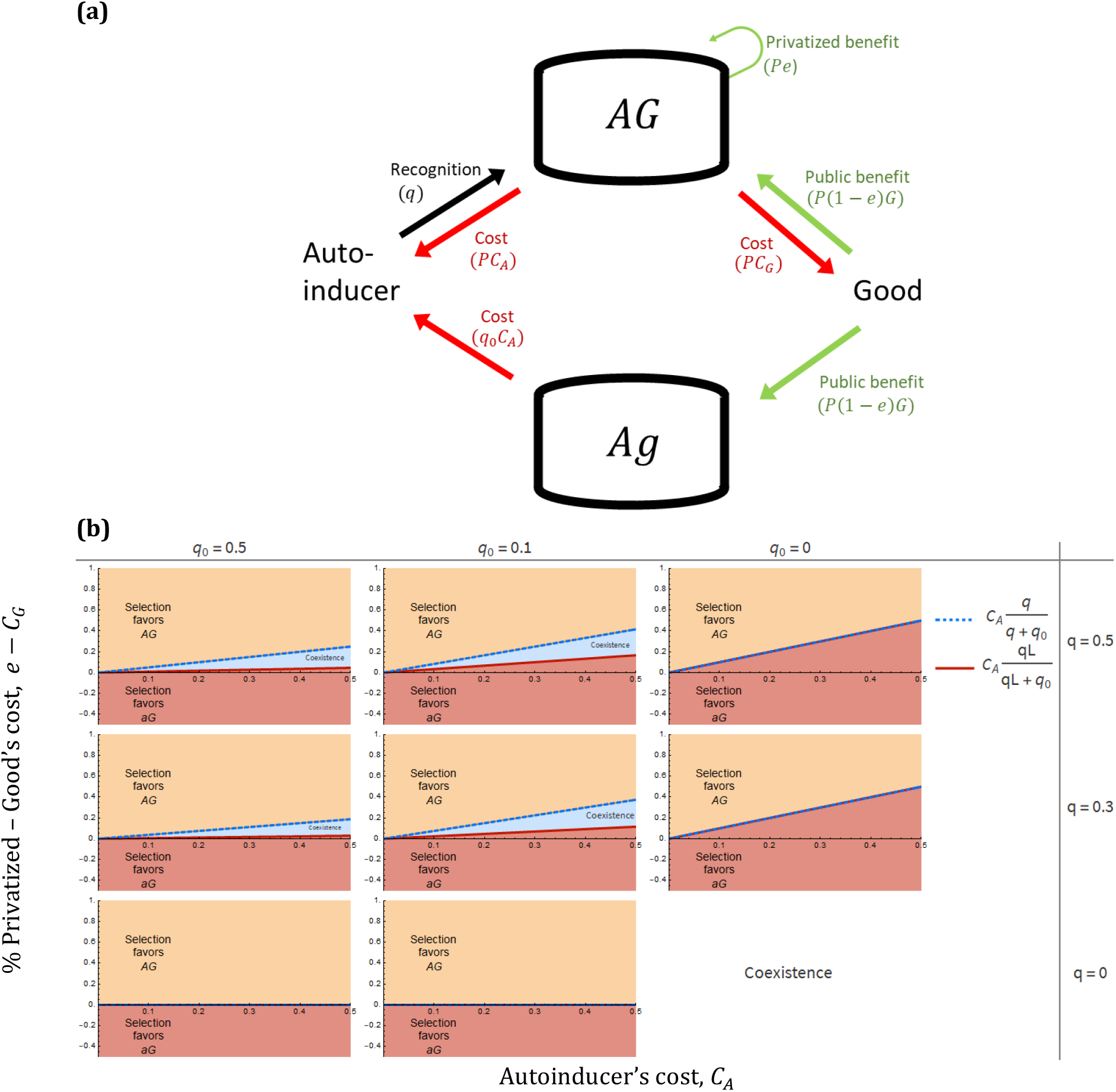
The partially privatized benefit fosters QS through pure populations of *AG* or through mixed populations of *AG* and *Ag*. **(a)** A schematic social interaction between strains *AG* and *Ag*. **(b)** Selection either favors (i) pure populations of *AG* (yellow area); (ii) pure populations of *Ag* (red area); (iii) coexistence of *AG* and *Ag* (blue area). Selection favors the coexistence if the rare strain outcompetes the common one (i.e., negative frequency-dependent selection). The frequency of each strain within the blue is not uniform (Figure S1). Negative frequency-dependent selection emerges from the co-regulation of QS (*q*) and QS-independent (*q*_*0*_) mechanisms, the existence of privatization (e) and having privatized benefits offsetting the minimum autoinducer’s cost, 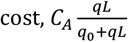, but not the maximum autoinducer’s cost, 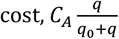. The x-axis is the per capita autoinducer’s cost, *C*_*A*_. The y-axis is the difference between the per capita partially privatized benefit and the per capita good’s cost, *e* − *C*_*G*_. Each subgraph captures the effect of regulatory architecture, via QS-dependent and QS-independent mechanisms. *q* = 0 implies absence of QS regulation. *q*_0_ = 0 implies absence of QS-independent regulation. Parameter: *L* = 0.1. Please see Table 2 and Section 6C for the definition of letters (q, C_A_ etc.) and the equations used for the analysis, respectively.

At any initial ratio of both strains, we found that selection can foster three outcomes. First, *AG* always outcompetes *Ag* if the difference between the total partially privatized benefit and the total good’s cost offsets the maximum autoinducer’s cost (yellow area in Figure 2b). Second, *Ag* always outcompetes *AG* if the difference between the total partially privatized benefit and the total good’s costs cannot offset the minimum autoinducer’s cost (red area in Figure 2b).

Lastly, selection fosters the coexistence of *AG* and *Ag* if the difference between the total partially privatized benefit and total good’s cost offsets the minimum autoinducer’s cost but not the maximum autoinducer’s cost. In this case, selection favors the rare strain over the common one (blue area in Figure 2b), i.e., negative frequency-dependent selection [18].

Moreover, we found that the ability to produce good in response to autoinducers does not stabilize the coexistence of AG and Ag.. This is because *AG* bears more autoinducer’s costs than *Ag* (Section 6.C). This is graphically noticeable in Figure 2B: the red area increases as the relative contribution of QS regulation increases (as *q* increases).

### (C) Pairwise analysis of *AG* and *aG*

As depicted in Figure 3, *aG* exploits *AG* by not producing autoinducers while use them to trigger the production of the good. Here, we analyzed whether QS can be maintained in a population by checking if selection fosters pure populations of *AG* or the coexistence of *AG* and *ag*.

**Figure 3.**
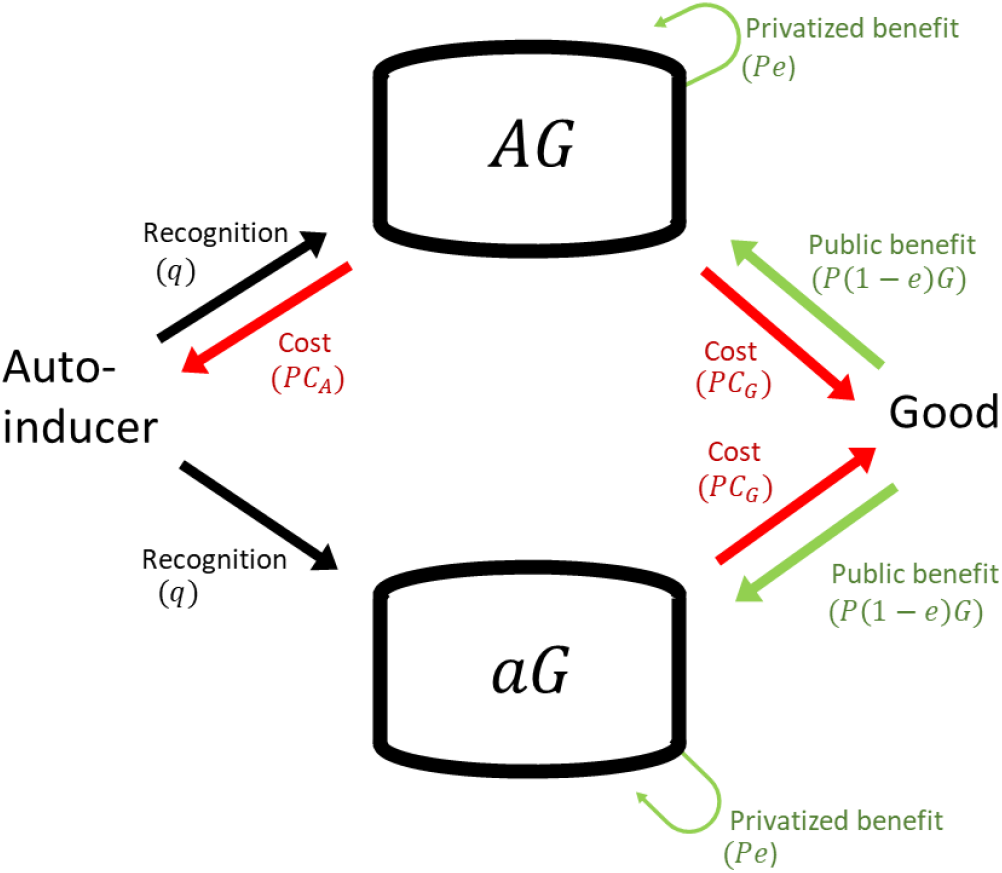
Partial privatization is ineffective against *aG* strains. Selection favors *aG* over *AG* because both strains access the private and public benefits and pay the good’s cost, but only *AG* pays the autoinducer’s cost. *aG* can use autoinducers to trigger the production of goods. *AG* can use autoinducers to trigger the production of more autoinducers and goods. Please see Table 2 and Section 6D for the definition of letters (q, C_A_ etc.) and the equations used for the analysis, respectively.

At any initial ratio of *AG* and *aG*, we found that selection always favors *aG* over *AG* (Equation 8). This is because both strains have the same access to benefits—including the partially privatized one—but only *AG* pays the autoinducer’s cost. This implies that the lower the per capita autoinducer’s cost, the more time is required for *aG* to eliminate *AG*.

### (D) Pairwise analysis of *Ag* and *aG*

As depicted in Figure 4, *Ag* exploits *aG* by not producing goods and yet accessing its public benefit. *aG* exploits *Ag* by not producing autoinducers and yet using them to trigger the production of goods. Hence, both strains have aligned interests in accessing each other’s molecules. Here, we analyzed whether QS can be maintained in a population by checking if selection fosters the coexistence of *AG* and *aG*.

**Figure 4.**
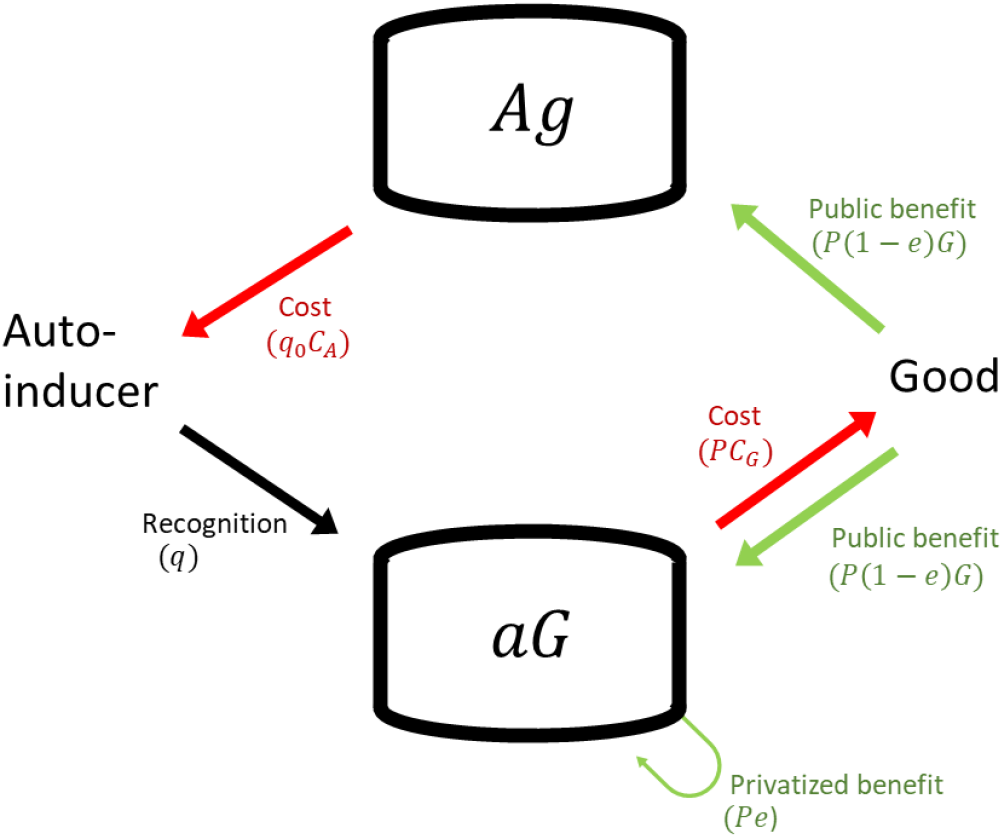
A schematic relationship for the social interaction between functionally complementary strains, i.e., the autoinducer-producer/good-nonproducer (*Ag*) and the autoinducer-nonproducer/good-producer (*aG*). Please see Table 2 and Section 6E for the definition of letters (q, C_A_ etc.) and the equations used for the analysis, respectively.

We found that selection cannot foster coexistence of *Ag* and *aG*. Thus, partial privatization cannot foster metabolic specialization, i.e., one strain produces good only and the other produces autoinducer only (Figure 5). Moreover, we found that if the per capita privatized benefit offsets the per capita good’s cost that is QS-regulated, *qL*(*C*_*G*_ − *e*) < 0, then *aG* always outcompetes *Ag* (Figure 5 c, f).

**Figure 5.**
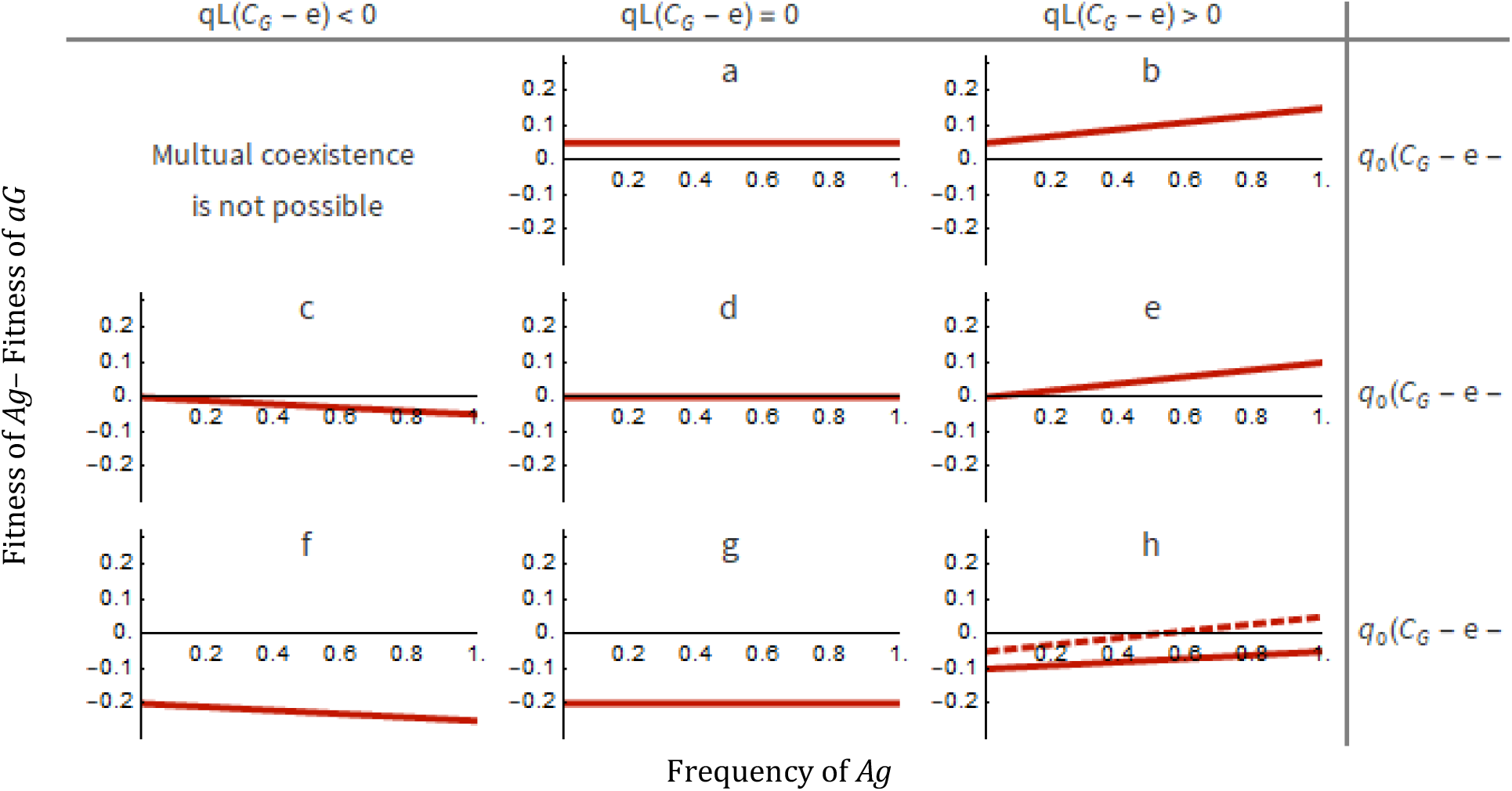
Partial privatization cannot foster coexistence of functionally complementary strains. The interaction between *Ag* and *aG* will always lead to pure populations of one of them. If the line is entirely below the x-axis, then *aG* always outcompetes *Ag*. If the line is entirely above the x-axis, then *Ag* always outcompetes *aG*. The point of intersection between the line and the x-axis indicates that both strains are equally fit. Selection cannot favor coexistence because the point of intersection only exists when the most common strain has the advantage over the rare strain (i.e., positive frequency dependence selection, dotted line). *qL*(*C*_*G*_ − *e*) is the inclination of the line and represents the metabolic balance from QS regulation. *q*_0_(*C*_*G*_ − *e* − *C*_*A*_) is the point of intersection with the y-axis and represents the metabolic balance from QS-independent regulation. Regarding the effect of QS regulation (*q*): *qL*(*C*_*G*_ − *e*) < 0 indicates that the per capita partially privatized benefit offsets the per capita good’s cost; *qL*(*C*_*G*_ − *e*) = 0 indicates a lack of QS regulation or a lack of QS-independent autoinducer secretion; *qL*(*C*_*G*_ − *e*) > 0 indicates that the per capita partially privatized benefit cannot offset the per capita good’s cost. Regarding the effect of QS-independent regulation (*q*_0_): *q*_0_(*C*_*G*_ − *e* − *C*_*A*_) > 0 indicates that the per capita good’s cost is higher than the sum of the per capita partially privatized benefit and the per capita autoinducer’s cost; *q*_0_(*C*_*G*_ − *e* − *C*_*A*_) = 0 indicates a lack of QS-independent regulation; *q*_0_(*C*_*G*_ − *e* − *C*_*A*_) < 0 indicates that the per capita good’s cost is lower than the sum of the per capita partially privatized benefit and the per capita autoinducer’s cost.

Otherwise, if the per capita privatized benefit does not offset the per capita good’s cost that is QS-regulated, *qL*(*C*_*G*_ − *e*) ≥ 0, then three outcomes are possible. First, *Ag* always outcompetes *aG* (line above x-axis). This will happen if the per capita good’s cost is higher than the sum of the per capita privatized benefit and the per capita autoinducer’s cost regulated by the QS-independent mechanism (Figure 5 A, B), i.e., *q*_0_(*C*_*G*_ − *e* − *C*_*A*_) > 0. *Ag* also always outcompetes *aG* if good’s synthesis is not regulated by the QS-independent mechanisms (Figure 5 E), i.e., *q*_0_(*C*_*G*_ − *e* − *C*_*A*_) = 0.

Second, *aG* always outcompetes *Ag* (line below x-axis). This will happen if the metabolic balance of QS-independent regulation is larger than the metabolic balance caused by QS-dependent regulation (Figure 5g, h, solid line), i.e., *q*_0_(*C*_*G*_ − *e* − *C*_*A*_) > *qL*(*C*_*G*_ − *e*). The metabolic balance of QS-independent regulation is the total difference between the per capita good’s cost and the sum of the per capita privatized benefit and the per capita autoinducer’s cost resulting from QS-independent regulation. The metabolic balance of QS regulation is the total difference between the per capita good’s cost and the per capita privatized benefit resulting from QS regulation.

Third, the common strain outcompetes the rare strain. This will happen if the metabolic balance of QS-independent regulation is smaller than the metabolic balance caused by QS-dependent regulation, i.e., *q*_0_(*C*_*G*_ − *e* − *C*_*A*_) < *qL*(*C*_*G*_ − *e*) (Figure 5h, dotted line).

### (E) Analysis of a population in which all the four strains exist

Above, we presented the analytical evolutionary outcome for pairwise interactions. However, the evolutionary dynamic in pairwise interactions need not be the same as when all strains are simultaneously interacting [16]. Here, we analyzed whether partial privatization fosters QS when all strains are simultaneously interacting (Figure 6a). To analyze whether QS can be maintained in a population, we examined if selection eliminates alleles *A* and *G* from the population, which would occur if *AG* were eliminated and if both *Ag* and *aG* were eliminated.

**Figure 6.**
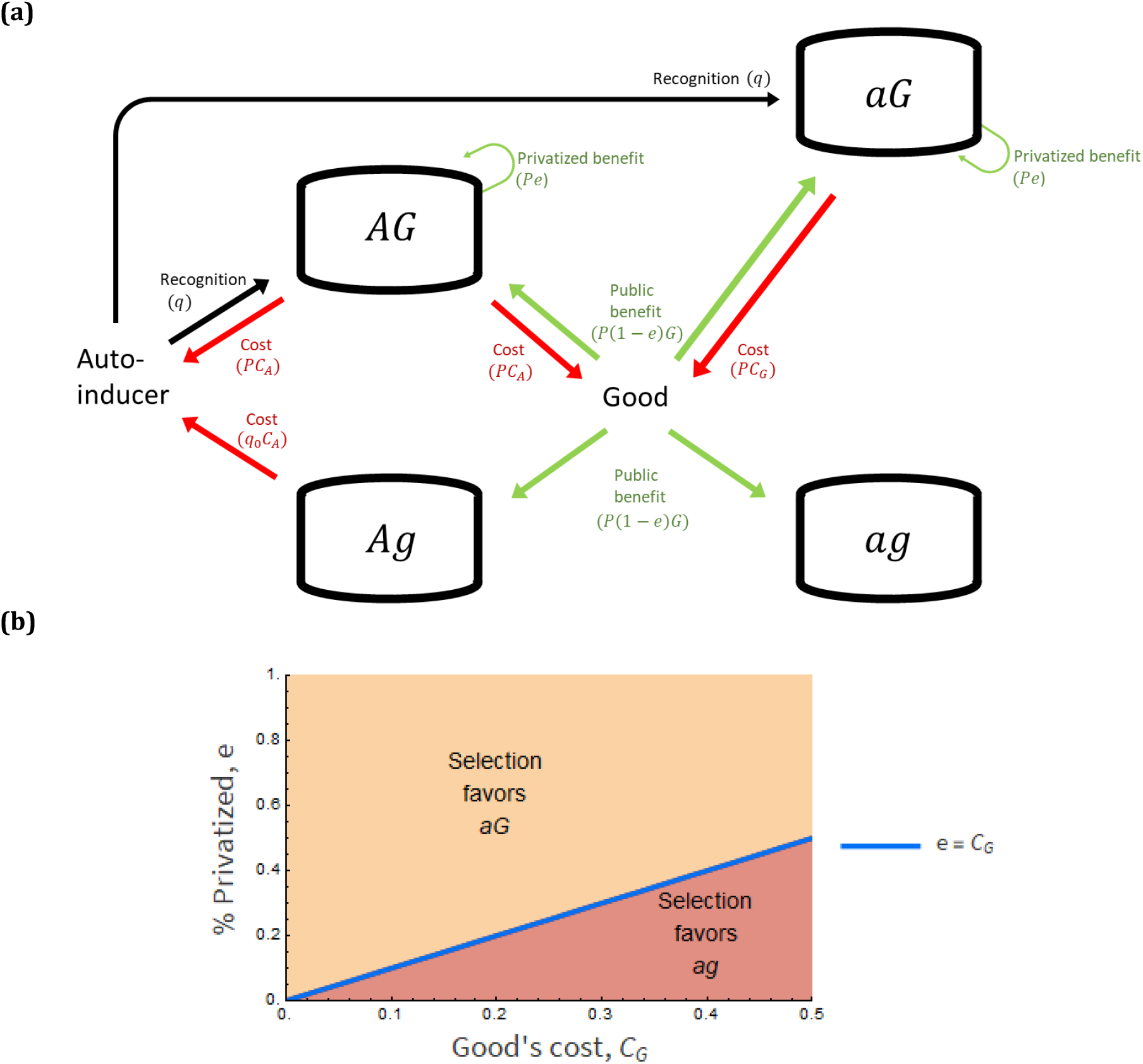
Partial privatization cannot foster QS when all strains are interacting. **(A)** A schematic relationship of when all strains are simultaneously interacting. **(B)** In an unstructured population, selection either favors pure populations of *aG* (yellow area) or pure populations of *ag* (red area). Selection always favors pure populations of *aG* if the per capita partially privatized benefit offsets the per capita good’s cost (*e* > *C*_*G*_). Otherwise (*e* < *C*_*G*_), selection always favors pure populations of *ag*. The blue line is when *aG* and *ag* are equally fit. This result holds for 0 ≤ *L* ≤ 1, 0 ≤ *q* ≤ 0.5, 0 ≤ *C*_*G*_ < 0.5, 0 ≤ *e* ≤ 1, 0 < *C*_*A*_ < 0.5, 0 < *q*_0_ < 0.5. Please see Table 2 and Section 6A for the definition of letters (q, C_A_ etc.) and the equations used for the analysis, respectively.

We found that partial privatization cannot foster QS. This is because selection leads to the loss of either allele *A* or both *A* and *G* alleles. Specifically, selection either fosters pure populations of *aG* or *ag* (Figure 6b). Pure populations of *aG* are favored if the per capita partially privatized benefit offsets the per capita good’s costs. Otherwise, selection favors pure populations of *ag*. Lastly, selection can only foster *AG* if autoinducers are costless, or if the *aG* strain is absent (SI).

After comparing these results with the pairwise interaction models, we noticed that pure populations of *AG*, pure populations of *Ag*, and mixed populations of *AG* and *Ag* are no longer favored by selection.

## 3. Discussion

Our work is the first to analyze the role of partial privatization in quorum sensing (QS) evolution. Our analytical results suggest that partial privatization of benefits cannot foster QS. This occurs because selection neither fosters (1) an *autoinducer-producer/good-producer* (fully producing, *AG*) strain nor (2) the coexistence of an *autoinducer-nonproducer/good-producer* (*aG*) and an *autoinducer-producer/good-nonproducer* (*Ag*), i.e., metabolic specialization of QS.

We found that partial privatization of good’s benefit cannot foster QS via a fully producing strain (Figure 6). This result occurs because both the fully producing strain (*AG*) and the *autoinducer-nonproducer/good-producer* strain (*aG*) access partially privatized benefits, but only the fully producing strain pays signaling costs (Equation 8). Thus, whenever the fully producing strain is interacting with *aG*, selection cannot foster the fully producing strain.

Although partial privatization is ineffective against the *aG* strain, partial privatization fosters the fully producing strain (*AG*) against the other two cheating strains (a*g* and *Ag*). This result occurs whenever the partially privatized benefit outweighs the cost of producing the mixed good and the autoinducer (Figure 1b, 2b). Additionally, our results show that negative frequency-dependent selection enables the coexistence between *AG* and *Ag* (blue area in Figure 2b, Figure S1). The negative frequency-dependent selection is known to stabilize coexistence in nature [18–21], which was also shown in some studies on QS evolution [22,23] and mixed good-producing strain’s evolution [5,12,14]. In *Pseudomonas aeruginosa*, experiments with the partial privatization of siderophores support our conclusion [7].

Here, we also found that partial privatization cannot foster QS (the coexistence of *A* and *G* alleles) via metabolic specialization. QS cannot be maintained by the coexistence between one strain specializing in autoinducer production (*Ag*) and one strain specializing in responding to it by producing a mixed good (*aG*) (Figure 5, S1). This prediction is in accordance with the absence of QS specialization in natural systems and experiments [22,24]. Also, our prediction agrees with the absence of a population that is mostly composed of *Ag-* and *aG*-like strains. [22,24]. However, previous studies on pairwise interactions [6,7,22,23,25–30] have not tested the interaction of *Ag-* and *aG*-like strains.

Why is metabolic specialization supported in previous models but not in ours [12,13,17]? There are two possible reasons that may explain this. First, while we have one mixed good and one autoinducer whose production is QS-dependent and QS-independent, earlier black queen models had two mixed goods being produced without the regulation by QS. Second, in earlier studies, metabolic specialization might occur because these models assumed that complementary strains are equally fit in their analysis [12,13,17]. While being equally fit is a justifiable assumption when each strain only produces one of two mixed goods, this assumption is unsuitable in our model because one strain only produces the autoinducer (*Ag*) and the other only the mixed good (*aG*). Assuming these complementary strains (i.e., *Ag* and *aG*) are equally fit is unsuitable because while mixed goods produce benefits directly, autoinducers indirectly produce benefits through the goods.

In our model, the effect of partial privatization was obtained by assuming linear benefits (the effect of benefits on bacteria is expressed as a linear function) and costs in an unstructured population. Therefore, a limitation of our model is that it does not include non-linear benefits/cost or the differential allocation of benefits towards genetically related individuals (i.e., kin selection). Earlier models (which assumed QS-independent regulation of mixed goods production) revealed coupling effects of nonlinear benefits from goods and kin selection. For instance, in yeast, while nonlinearity of benefits from sucrose metabolism explains selection favoring the coexistence between cooperators and cheaters, linearity of benefits does not [3]. Moreover, the coupled effect of partial privatization and spatial allocation of goods favored cooperation more than partial privatization alone [12].

Additionally, earlier QS models (which assumed no partial privatization) revealed that nonlinearity [31–33] and kin selection [34–36] dramatically influence the direction and strength of selection. Thus, our result that an *aG* always outcompetes *AG* (Section 6D) might change if we consider the coupled effect of partial privatization and kin selection. It would be of interest to study QS evolution in our model incorporating nonlinearity and kin selection.

## 4. Conclusion

Our analytical results show how physiological partial privatization of benefits might affect QS evolution. We have shown that partial privatization cannot explain why selection favors QS in a large unstructured population. Our findings suggest that, if the partially privatized benefit cannot offset the production costs, the evolution of QS in a bacterial population with partial privatization is qualitatively similar to the cases without partial privatization. On the other hand, if the partially privatized benefit offsets the production costs, the evolution of QS in a bacterial population with partial privatization is not reducible to the cases without partial privatization. Thus, our study indicated that new studies are needed to evaluate the potential underrepresentation of partial privatization on QS evolution.

## 5. Model

### Model framework

We consider a large clonally reproducing population, large enough that the probability of a loss of a temporarily rare gene by random fluctuations is negligible. The population has discrete non-overlapping generations. To single out the effect of partial privatization, we assume that autoinducers and mixed goods are homogenously distributed across the environment at all times. We also assume that social interactions occur in an unstructured population; that is, cells are homogeneously distributed and there is no migration.

Using the standard population genetic framework [16], we track the change in strains’ frequency through time

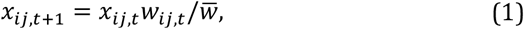

where, *x*_*ij,t*_ is the frequency of strain *ij* at time *t, i* = {*A, a*} and *j* = {*G, g*}. *w*_*ij*_ is the fitness of strain *ij* at time *t*. 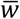 is the population mean fitness. Below we describe strains’ behavior and fitness.

### Strains and fitnesses

We consider two types of secreted molecules. One is a costly autoinducer. The other is a costly mixed good. Mixed goods can produce benefits by promoting growth and by reducing mortality. Here, we assumed that the mixed good promotes growth. We assume that there is no external source of autoinducers and mixed goods, i.e., these two molecules are only biologically produced.

In natural systems, the more autoinducers are present in the environment, the higher the secretion of autoinducers (positive feedback loop). We indirectly modeled this positive feedback loop by assuming that *AG* always secretes more autoinducer than *Ag*. We did this by assuming that both *AG* and *Ag* secrete a QS-independent fraction of autoinducers, *L*, but only *AG* produces the fraction of autoinducers regulated by QS-dependent regulation, (1 − *L*). Thus, the frequency of autoinducers in the environment at time *t, A*_*t*_, is the sum of QS-independent, *L* (*x*_*AG,t*_ + *x*_*Ag,t*_), and QS-dependent, (1 − *L*)*x*_*AG,t*_ production of autoinducer

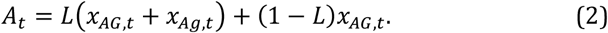

We assume that the production of the autoinducer has a per capita cost *C*_*A*_, 0 ≤ *C*_*A*_ < 0.5. The production of the good has a per capita cost *C*_*G*_, 0 ≤ *C*_*G*_ < 0.5. The total cost paid is positively correlated to an autoinducer’s frequency. Let *qA*_*t*_ be the probability of gene activation given an autoinducer’s frequency. Let *q* be the constant rate at which an autoinducer and a good are produced given *A*_*t*_. That is, *q* represents the influence of QS regulation, the ability of kin discrimination. Based on empirical findings that some QS-regulated mixed goods are co-regulated by QS-independent mechanisms [37,38], we consider the constant rate *q*_0_. Thus, the probability of an autoinducer’s and a good’s production is

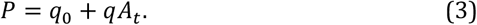

Given that 0 ≤ *A*_*t*_ ≤ 1, we ensure that *P* varies between 0 and 1, by having 0 ≤ *q*_0_, *q* ≤ 1/2. *q*_0_ = 0 indicates the lack of QS-independent regulation. *q* = 0 indicates absence of QS regulation. We allow *q*_0_ to be larger than *q* because research found that the regulatory genetic architecture can evolve relatively fast to become less reliant on QS [39].

Only *AG* and *Ag* pay an autoinducer’s cost. The total cost for autoinducer’s production for *Ag* is *q*_0_*C*_*A*_. Because *Ag* cannot recognize autoinducers, its regulation only occurs via a QS-independent mechanism. The total cost for autoinducer’s production for *AG* is *PC*_*A*_. Because *AG* can recognize autoinducers, its regulation occurs via QS and QS-independent mechanisms. Only *AG* and *aG* produce the mixed good. *AG* and *aG* pay the cost of goods regulated and not regulated by QS, *PC*_*G*_.

The frequency of goods in the environment is *PG*_*t*_. *G*_*t*_ is the frequency of allele *G* at time *t*, i.e., *G*_*t*_ = *x*_*AG,t*_ + *x*_*aG,t*_. The benefit generated by allele *G* is unity. Let *e* (0 ≤ *e* ≤ 1) be the fraction of the benefit that is of exclusive access to producers, i.e., the percentage of partially privatized benefit. *e* = 0 implies that the whole benefit is shared, i.e., fully public. *e* = 1 implies that the whole benefit is private. (1 − *e*) is the fraction of benefit that is public. Thus, 1*e* and 1(1 − *e*)*G* are the per capita private and the per capita public benefit, respectively. *Pe* and *P*(1 − *e*)*G* are the total private and the total public benefit, respectively. We assume a baseline fitness of unity. Taking together, the fitnesses equations are

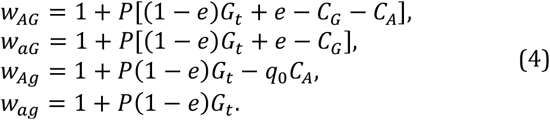

A list of parameters and variables are found in Table 2.

## 6. Analytical solutions

### (A) All strains simultaneously interacting

Let *A*_*t*_ = *L*(*x*_*AG,t*_ + *x*_*Ag,t*_) + (1 − *L*)*x*_*AG,t*_ and *G*_*t*_ = *x*_*AG,t*_ + *x*_*aG,t*_ be the autoinducer’s and good’s frequency in the environment at time *t*. By eliminating the common terms of Equation 4, we have the simplified fitness equations

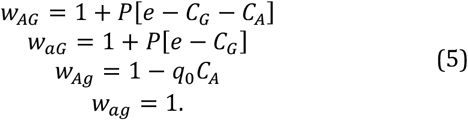

From the fitness equations above, it is clear that *w*_*aG*_ ≥ *w*_*AG*_ and *w*_*ag*_ ≥ *w*_*Ag*_. We found that selection will either drive evolution towards pure populations of *aG* or *ag*. Selection favors pure populations of *aG* (*x*_*aG*_ = 1) if *e* > *C*_*G*_. Otherwise (*e* < *C*_*G*_), selection favors pure populations of *ag* (*x*_*ag*_ = 1). Selection only stops favoring these outcomes if *q*_0_ = 0 or *C*_*A*_ = 0. Under these parameter values, *w*_*AG*_ = *w*_*aG*_ and *w*_*Ag*_ = *w*_*ag*_.

### (B) Pairwise interaction between *AG* and *ag* strains

Let *A*_*t*_ = *x*_*AG,t*_ be the autoinducer’s frequency at time *t*. Let *G*_*t*_ = *x*_*AG,t*_ be the maximum good’s frequency in the environment at time *t*. The relative fitness of *AG* (*w*_*AG*_ > *w*_*ag*_) is

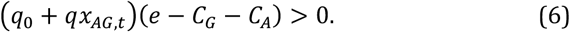

Selection favors pure populations of *AG* (*x*_*AG*_ = 1) if *e* > *C*_*G*_ + *C*_*A*_, *q*_0_ ≠ 0 and *q* ≠ 0. Selection favors pure populations of *ag* (*x*_*AG*_ = 1) if *e* < *C*_*G*_ + *C*_*A*_ and *q*_0_ ≠ 0.

### (C) Pairwise interaction between AG and *Ag* strains

Let *A*_*t*_ = *L* + (1 − *L*)*x*_*AG,t*_ be the autoinducer’s frequency at time *t*. Let *G*_*t*_ = *x*_*AG,t*_ be the maximum good’s frequency at time *t*. The relative fitness of *AG* (*w*_*AG*_ > *w*_*Ag*_) is

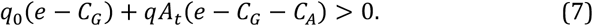

Selection can foster three possible outcomes. First, a pure population of *AG* (*x* _*AG*_ = 1) is stable if 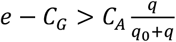. Second, a pure population of *Ag* (*x*_*Ag*_ = 1) is stable if 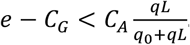 Lastly, a mixed population of *AG* and 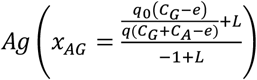 is stable if 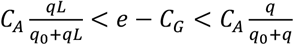.

### (D) Pairwise interaction between *AG* and *aG* strains

Let *A*_*t*_ = *x*_*AG,t*_ be the autoinducer’s frequency at time *t*. Let *G*_*t*_ = *x*_*AG,t*_ + *x*_*aG,t*_ = 1 be the good’s frequency at time *t*. The difference in fitness between *AG* and *aG* is

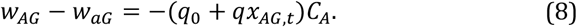

Thus, selection always favors pure populations of *aG* (*x*_*aG*_ = 1), unless the production of autoinducer is costless (*C*_*A*_ = 0) or the regulatory mechanisms become defective (*q*_0_ = 0 and *q* = 0), which implies that both strains are equally fit.

### (E) Pairwise interaction between functionally complementary strains (i.e., *Ag*, and *aG*)

Let *A*_*t*_ = *Lx*_*Ag,t*_ be the autoinducer’s frequency at time *t*. Let *G*_*t*_ = *x*_*aG,t*_ be the maximum good’s frequency at time *t*. The relative fitness of *Ag* (*w*_*Ag*_ > *w*_*aG*_) is

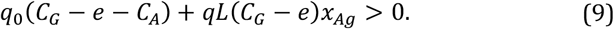

We found that if the per capita private benefit offsets the per capita good’s cost (*C*_*G*_ < *e*), then *aG* always outcompetes *Ag* (i.e., *w*_*aG*_ > *w*_*Ag*_ is always true) (Figure 5 c, f). Otherwise (*C*_*G*_ > *e*), selection can generate one of three possible outcomes.

First, *Ag* always outcompetes *aG* (Figure 5 b,e). This happens if *qL*(*C*_*G*_ − *e*) ≥ 0 and *q*_0_(*C*_*G*_ − *e* − *C*_*A*_) ≥ 0. Meaning, if the autoinducer cost paid by *Ag* is not enough to cause the net deficit in producing goods (*C*_*G*_ > *e*) paid by *aG*.

Second, *aG* always outcompetes *Ag* (Figure 5h, solid line). This happens if *qL*(*C*_*G*_ − *e*) ≥ 0, *q*_0_(*C*_*G*_ − e − C_A_) < 0 and *qL*(*C*_*G*_ − *e*) < *q*_0_(*C*_*G*_ − *e* − *C*_*A*_). That is, if the difference between the per capita good’s cost is lower than the sum of the per capita privatized benefit and the per capita autoinducer’s cost and if the total of this metabolic balance caused by QS-independent regulation, *q*_0_(*C*_*G*_ − e − C_A_) < 0, is larger than the metabolic balance coming from the QS-regulation generating a per capita good’s cost and a per capita privatized benefit, *qL*(*C*_*G*_ − *e*) < *q*_0_(*C*_*G*_ − *e* − *C*_*A*_).

Third, positive frequency-dependent selection favors the most common strain (Figure 5h, dotted line). This happens if *qL*(*C*_*G*_ − *e*) ≥ 0, *q*_0_(*C*_*G*_ − e − C_A_) < 0 and *qL*(*C*_*G*_ − *e*) > *q*_0_(*C*_*G*_ − *e* − *C*_*A*_). That is, if the metabolic balance of QS-independent regulation is smaller than the metabolic balance caused by QS regulation.

## Supplemental information

**Figure S1.**
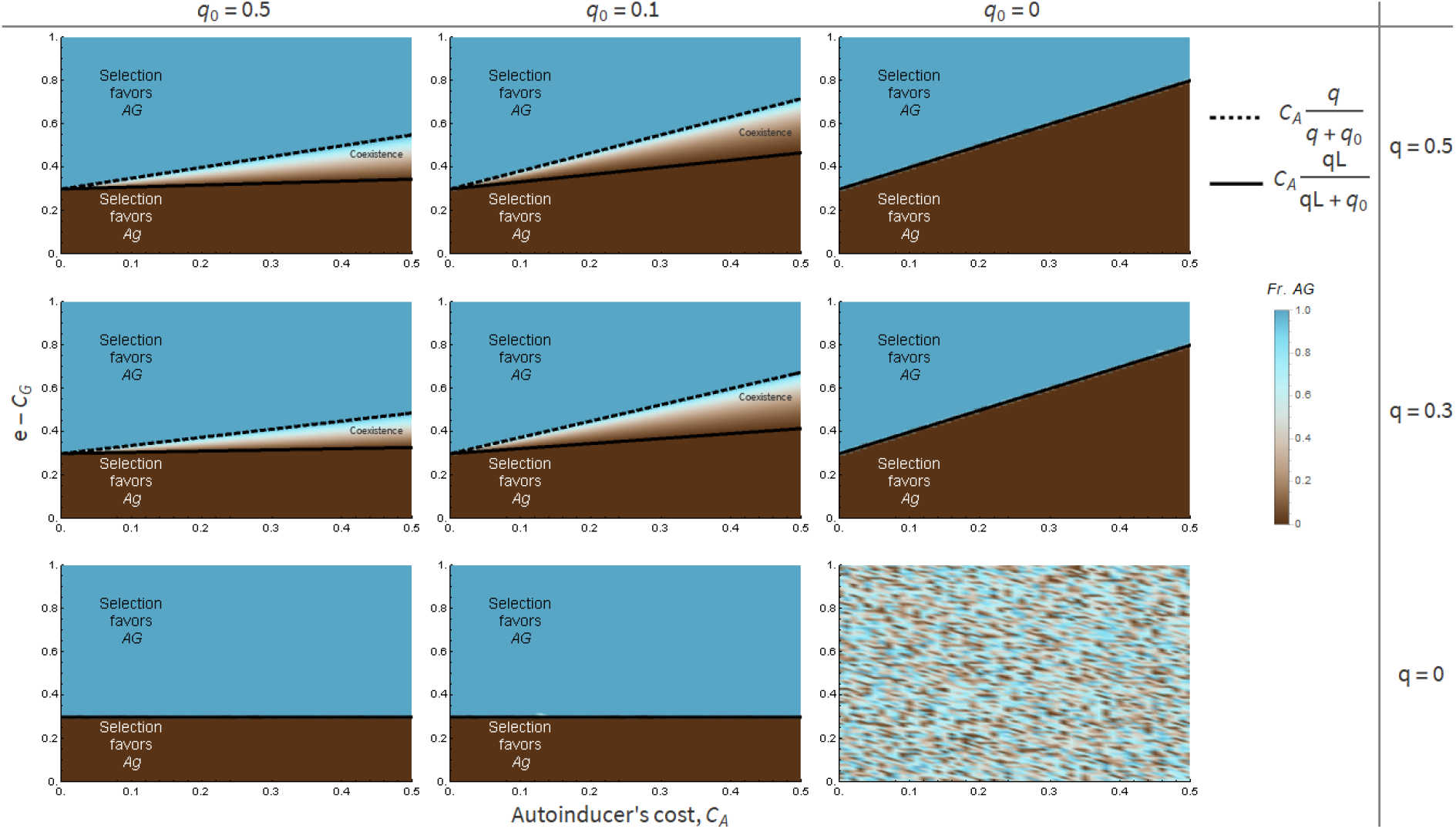
The partially privatized benefit fosters QS through pure populations of *AG* or through mixed populations of *AG* and *Ag*. Selection either favors (i) pure populations of *AG* (blue area); (ii) pure populations of *Ag* (brown area); (iii) coexistence of *AG* and *Ag*. Selection favors the coexistence if the rare strain outcompetes the common one (i.e., negative frequency-dependent selection). Negative frequency-dependent selection emerges from the co-regulation of QS (q) and QS-independent (q_0_) mechanisms, the existence of privatization (e) and having privatized benefits offsetting the minimum autoinducer’s cost, 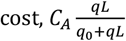, but not the maximum autoinducer’s cost,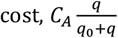. The x-axis is the per capita autoinducer’s cost, *C*_*A*_. The y-axis is the difference between the per capita partially privatized benefit and the per capita good’s cost, *e* − *C*_*G*_. Each subgraph captures the effect of regulatory architecture, via QS-dependent and QS-independent mechanisms. *q* = 0 implies absence of QS regulation. *q*_0_ = 0 implies absence of QS-independent regulation. Parameters: *C*_*G*_ = 0.3, *L* = 0.1. Initial frequency of each strain was draw from a uniform distribution. The simulation was stopped after 10000 steps.

